# Population stratification in GWAS meta-analysis should be standardized to the best available reference datasets

**DOI:** 10.1101/2020.09.03.281568

**Authors:** Aliya Sarmanova, Tim Morris, Daniel John Lawson

## Abstract

Population stratification has recently been demonstrated to bias genetic studies even in relatively homogeneous populations such as within the British Isles. A key component to correcting for stratification in genome-wide association studies (GWAS) is accurately identifying and controlling for the underlying structure present in the sample. Meta-analysis across cohorts is increasingly important for achieving very large sample sizes, but comes with the major disadvantage that each individual cohort corrects for different population stratification. Here we demonstrate that correcting for structure against an external reference adds significant value to meta-analysis. We treat the UK Biobank as a collection of smaller studies, each of which is geographically localised. We provide software to standardize an external dataset against a reference, provide the UK Biobank principal component loadings for this purpose, and demonstrate the value of this with an analysis of the geographically sampled ALSPAC cohort.

## Introduction

Genome-wide association studies (GWAS) are increasingly being used to identify biological pathways underlying complex traits and diseases. They have become an essential part of making direct links between genetics and phenotypes (Visscher et al. 2017) and have facilitated causal inference through Mendelian Randomization (Paternoster, Tilling, and Smith 2017; Zhu et al. 2018). However, detecting and interpreting associations remains a challenge because genetic associations tend to be tiny (particularly for polygenic traits) and other associations may be large.

Many groups have joined efforts to create large consortia that assemble results from multiple GWAS, providing aggregated sample sizes that are now in excess of a million individuals (Linnér et al. 2019; Lee et al. 2018). Meta-analysis of consortia datasets improves the power necessary to detect many genotype-phenotype associations. However, where population structure exists in a dataset but is insufficiently controlled for, it can lead to spurious or inflated genotype-phenotype associations (Lawson et al. 2020; Peterson et al. 2017). Even within the UK, considering only white people of European ancestry, migration and socio-economic position correlate with ancestry (Abdellaoui et al. 2019; Haworth et al. 2019).

Recently it has become apparent that GWAS results based on large scale meta-analysis have been at least partially biased due to inadequate correction for confounding by population stratification. The Genetic Investigation of ANthropometric Traits (GIANT Consortium 2018) meta-analysis of height and BMI (Wood et al. 2014; Locke et al. 2015) has led to ambiguous conclusions regarding selection on height (Yengo et al. 2018)1 with Genetic Scores being of particularly discussion vulnerable to this confounding (Berg et al. 2019; Sohail et al. 2019). Similar issues have been reported for Educational Attainment (Abdellaoui et al. 2019; Haworth et al. 2019), as well as and diseases including Type 2 diabetes and coronary heart disease (Reisberg et al. 2017).

Latent structure and population stratification are addressed during the discovery of associated genetic variants by correcting for Principal Components (PCs) of the genetic variation (Price et al. 2006). Historically, only a few PCs were used, increasing with sample size and time from two (Wellcome Trust Case Control Consortium 2007), five (Warrington et al. 2015) and ten (Okbay et al. 2016) to 40 - the default provided by UK Biobank (Bycroft et al. 2018) – to 100 or more (Abdellaoui et al. 2019). Yet even 100 PCs are insufficient (Lawson et al. 2020) as important structures may explain less variation than noise and hence remain uncorrected, which can lead to uneven correction and bias in meta-analysis.

We propose a simple solution. Correction can be improved and standardized using a large external reference dataset to define “all human genetic variation”, against which local variation within a single study can be compared. Thus, whilst population stratification might act as a source of covariance between genotypes and phenotypes, this can be corrected for. We demonstrate that meta-analyses corrected for population stratification using a large external reference dataset (“global” ancestry correction) performs better than meta-analyses corrected for population stratification using the same dataset (“local” ancestry) in the UK Biobank.

There are alternative methodologies for stratification correction that go beyond PC correction. Linear Mixed Models (LMMs) have gained popularity since their introduction in genetics (Yu et al. 2006) through easy-to-use software such as GCTA (Yang et al. 2011). LMMs are a “gold-standard” for GWAS because instead of correcting for only the top few variance components, they correct in principle for the entire Genetic Relatedness Matrix (GRM) comparing all pairs of individuals. This allows familial structure to be corrected in the same framework as ancestry. However, whilst pairwise relationships in the GRM are measured in the data, correlations between them are still estimated with noise and hence correction performance improves with sample size. We use Bolt-LMM (Loh et al. 2015) throughout and find that correction for external PCs complements, and is not replaced by, the use of LMMs.

We investigate the relationship between latent genetic structure and phenotypes, i.e. population stratification, in the UK Biobank. We demonstrate that proper correction for stratification has implications in the Avon Longitudinal Study of Parents and Children (ALSPAC) (Boyd et al. 2013; Fraser et al. 2013) in Bristol, UK, especially were the results are to be considered as part of a meta-analysis. These findings provide evidence that similar correction will lead to changes in findings for large-scale meta-analysis.

Software and appropriate reference data are provided (see Code Availability) to allow others to easily apply this to their own data.

## Results

### Identification of population structure required for correction

Successful identification and prioritization of disease-associated causal variants relies on understanding the distribution of genetic variants within and between populations. However, the extent to which ancestry can impact variant frequencies is not always clear. Accurate understanding and use of methods of correcting for ancestry such as PCs is critical.

We are interested in constructing and improving ancestry inference for all studies. To this aim we constructed 200 PCs (see Materials and Methods) following the sample and SNP selection and PC computation methodology of (Bycroft et al. 2018). Critically, PC loadings and eigenvalues are made available, allowing projection of external datasets into this ancestry measure, which we refer to as “global” ancestry/PCs. This contrasts to “local” ancestry and PCs, constructed using PC analysis within a single dataset.

The global moniker implies usefulness outside of the UK. The UK Biobank remains one of the largest easily accessed resources for worldwide variation, including (with some arbitrary choices of definition) over 6k Sub-Saharan Africans, 2k East Asians, and 7k South Asians. Naturally, a larger reference would identify further local structure. Similar to a recent study (Privé et al. 2020), we found evidence (Supplementary Figure S1) that Linkage Disequilibrium (LD) is important after the first 18 PCs, that ancestry associations reduce after 40 PCs, and that some population structure is associated with further PCs (Materials and Methods).

### Population Structure in the UK Biobank

We restricted our stratification analyses to 331,890 UK Biobank participants of UK ancestry excluding Northern Ireland, and ∼12M SNPs after quality-control filtering and LD pruning (see Materials and Methods). For illustration purposes, we clustered individuals using k-means (see Materials and Methods) into 5 clusters (Figure 1a). The largest cluster represented southern and eastern England, with northern England, Scotland, North Wales, and South Wales each being represented (Galinsky et al. 2016). We are not attempting to infer actual ancestry from these PCs.

**Figure 1.**
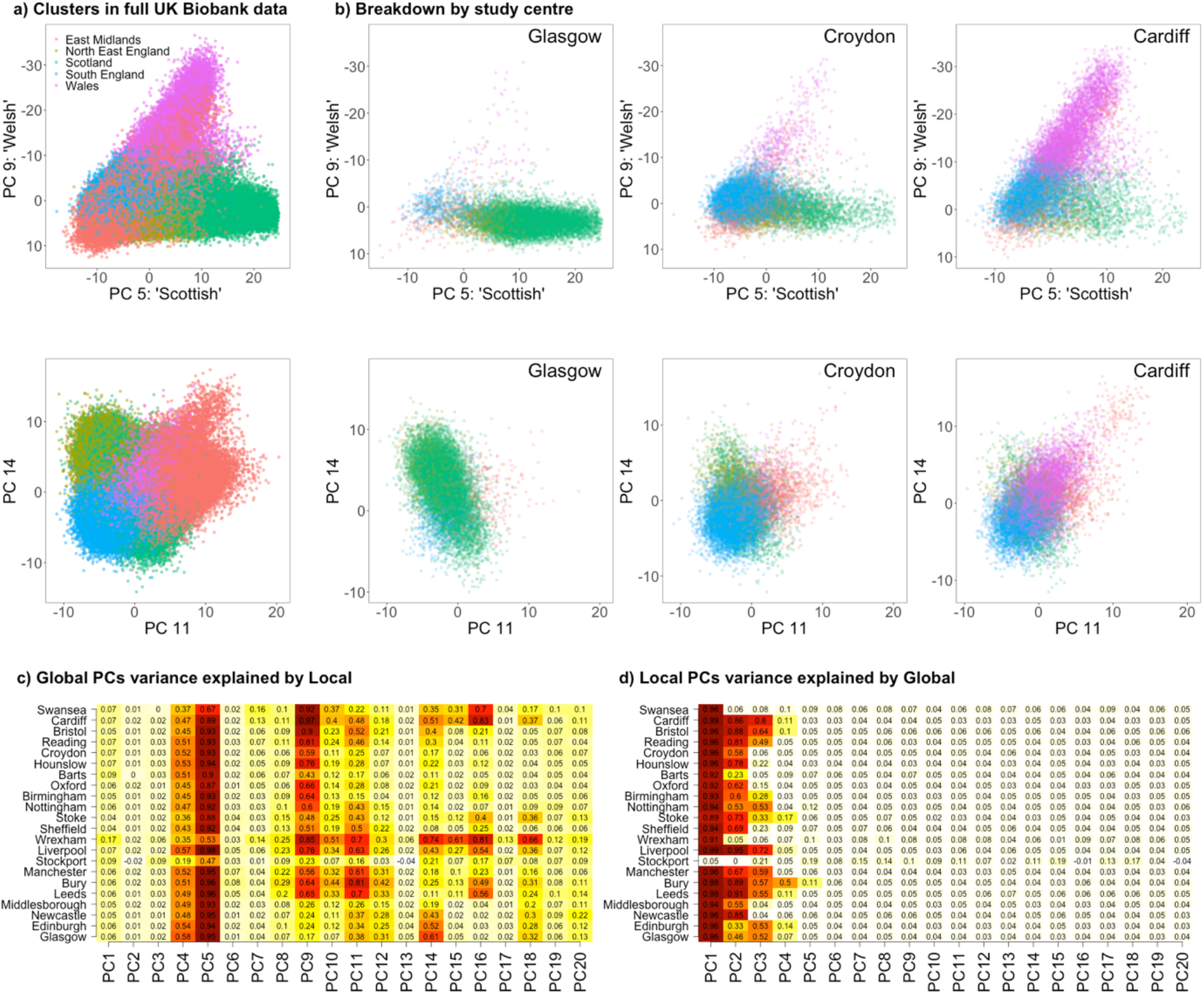
UK Biobank PCs by study centre Global. (i.e. inferred in the full UK Biobank) genetic ancestry PCs (Principal Components) is incompletely captured by local ancestry. a) The Global (whole biobank) PC analysis reveals British ancestry primarily in PCs 5,9,11 and 14 (see Supplementary Figure S2). b) Retaining PCs only for one geographical study centre at a time shows that many ancestries are under-sampled. c) Conducting a PC analysis within a single study centre, and trying to recover the PCs (see Methods), leads to low variance explained (R^2^) for many PCs. d) Predicting in reverse, only the first 2-5 PCs of a local analysis capture ancestry, with the remaining PCs being non-significant and are shown in pale with a white border (see Methods).

PCs are ordered by the total variation explained in the data. Major variation directions are associated with deep historical splits between populations such as African vs Eurasians (PC1-2), Europeans vs East Asians (PC1-3), Central Asia (PC3-4), and Europe (PCs 5,8). This contrasts regional variation within the UK for which the main PCs are 5 and 9 describing variation between English, Scottish and Welsh ancestry, as we as PCs 11 and 14 which further separate structure within Wales and England. This is strongly structured by study centre, which captures current living location (Figure 1b). These and other PCs (Supplementary Figure S2-3) correspond to known historical and geographical areas (Leslie et al. 2015).

To assess how much of this variation is captured by local PCs, we performed PC projection, i.e. a regression analysis for each global PC using all local PCs as predictors (see Materials and Methods). Local PCs capture global variation with varying veracity (Figure 1c). The predictability of global PCs varies by study centre according to which populations are poorly represented in them. PC5 is best explained in the West and describes Welsh vs English ancestry. PC9 describes South Wales ancestry; PC11 describes northern England ancestry; PC14 describes Scottish ancestry; whilst PC16 describes North Welsh ancestry. Worldwide ancestry PCs are homogeneous within the UK and therefore cannot be explained (PC1-3,6-8,13,17). Local PCs for all 22 study centres fail to explain some UK ancestry, and the inverse prediction of explaining local PCs using Global PCs shows that the local analyses typically contain only 2-4 ancestry related PCs (Figure 1d).

This observed population structure within the UK provides a source of covariance between genotypes and phenotypes that can bias epidemiological inference from genetic data. The following sections establish consequences of unexplained covariance for understanding complex disease.

### Stratification correction using global vs local PCs in UK Biobank

The most straightforward measure of stratification is of the total variation in phenotypes explained by genetic PCs, without attributing this to individual SNPs. Both educational attainment (EA) and Body Mass Index (BMI) vary by region (Supplementary Figure S4) and show large systematic differences between local ancestry and global ancestry correction (Figure 2). Several study centres explain dramatically less variation with local PCs than global, for example for EA in Croydon (0.6% local vs 3.2% global) and Hounslow (0.8% local vs 3% global). Figure 1c-d explains this as a failure to identify components corresponding to Scottish, Welsh and other ancestries that are individually rare but nevertheless important when considered together. Conversely others, especially centres with small sample sizes such as Wrexham and Swansea, explain more variation in local than global ancestry.

**Figure 2.**
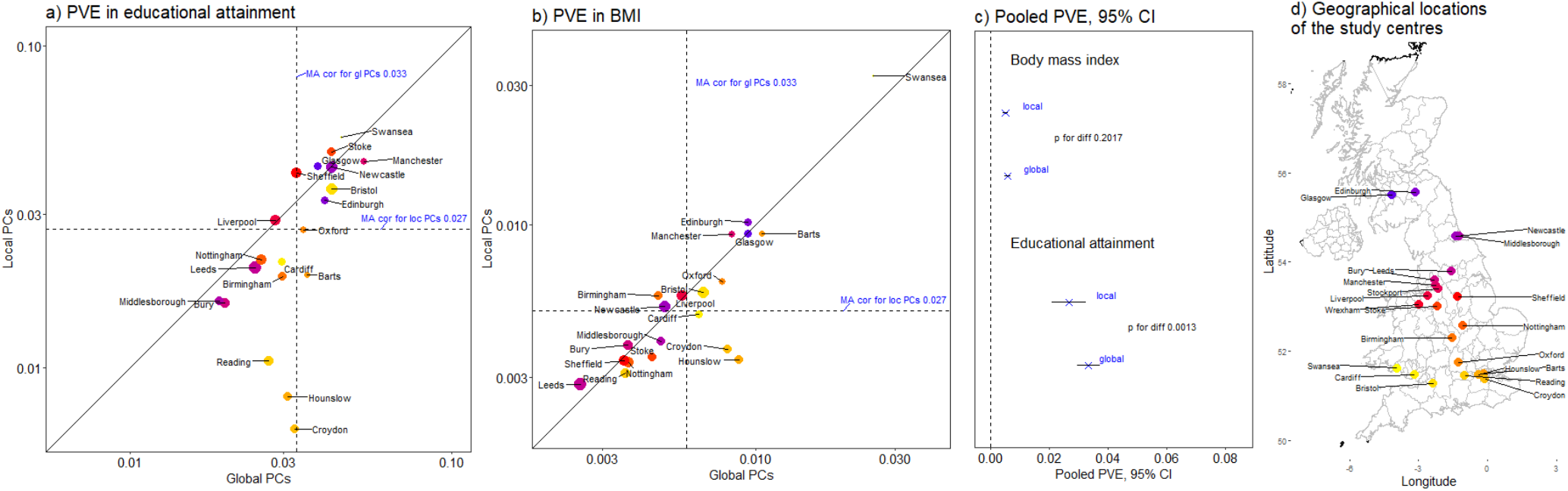
Stratification correction bias seen in Proportion of Variance Explained (PVE) Meta-analysis of UK Biobank study centres demonstrates stratification problems. a) Proportion of Variance Explained (PVE) in Educational attainment corrected for 40 global vs 40 local PCs, split by study centre. The point size indicate sample size per study centre, and colours show geography (d). b) Proportion of Variance Explained in BMI (Body Mass Index). c) Pooled PVE and 95% Confidence Intervals, with p-values for a paired t-test for a difference in mean. d) The Geographical locations of the study centres explaining the colour gradient: from Scotland/North (blue) to Midlands (red), via South-East (orange) to Wales/South-West (yellow).

We tested 24 disease statuses for the amount of variance explained by Local or Global PC correction, and found that Psoriasis, Hyperthyroidism, and Hypothyroidism were all significant different (Figure S5) and Multiple Sclerosis and Asthma are implicated though not significant after correcting for multiple testing.

Our analyses demonstrate two competing effects. Firstly, local PCs in small studies “overfit”, as they are able to explain much of the variance present regardless of whether it describes real ancestry or noise. This is why the number of PCs corrected for is often thresholded using a noise-level approximation (Lawson and Falush 2012) and justifies the small number of PCs used in early GWAS. Secondly, some ancestry components will not be recovered in a small dataset due to lack of statistical power. Mathematically, PC analysis displays a transition as sample size decreases, in which a particular population structure is identified when enough variation exists for it, and rather abruptly becomes indistinguishable from noise (McVean 2009). Importantly, local PCs perform worse not solely in small studies, but in larger but genetically more homogenous populations of the South-East of England. It is rare shared variation, regardless of the size of the study, that local PCs fail to identify and hence correct for.

### Local vs global correction for individual GWAS Effect sizes in UK Biobank

Meta-analysis is a statistical tool for combining results from coherent studies on different samples. A fundamental principle in GWAS meta-analysis is that all studies included examined the same hypothesis, had similar study design and analyzed study-level SNPs in a near-identical way (Zeggini and Ioannidis 2009; Bush and Moore 2012; Evangelou and Ioannidis 2013), similar imputation (Li et al. 2009), quality control, large-scale ancestry (Peterson et al. 2017) and of course, population stratification correction. Meta-analysis is individually important and offers a chance to examine stratification correction entirely within the (supposedly) homogeneous UK Biobank cohort.

For EA and BMI we estimated effect sizes when performing meta-analysis with global and local PC correction in the UK Biobank. Whilst individually, most SNP effect changes are not statistically significant, three issues arise (Figure 3). Firstly, estimates are systematically larger in magnitude when correcting with local rather than global ancestry. Secondly, some subsets of SNPs respond in a systematically different way (Supplementary Figure S6), leading to “clusters” of SNPs that are under, or over, corrected using local ancestry alone. Finally, smaller effects with the least statistical support are larger with local correction; by 2% in EA, 0.6% for BMI for genome-wide significant SNPs (determined by regressing local estimates on global; Supplementary Figure S6).

**Figure 3.**
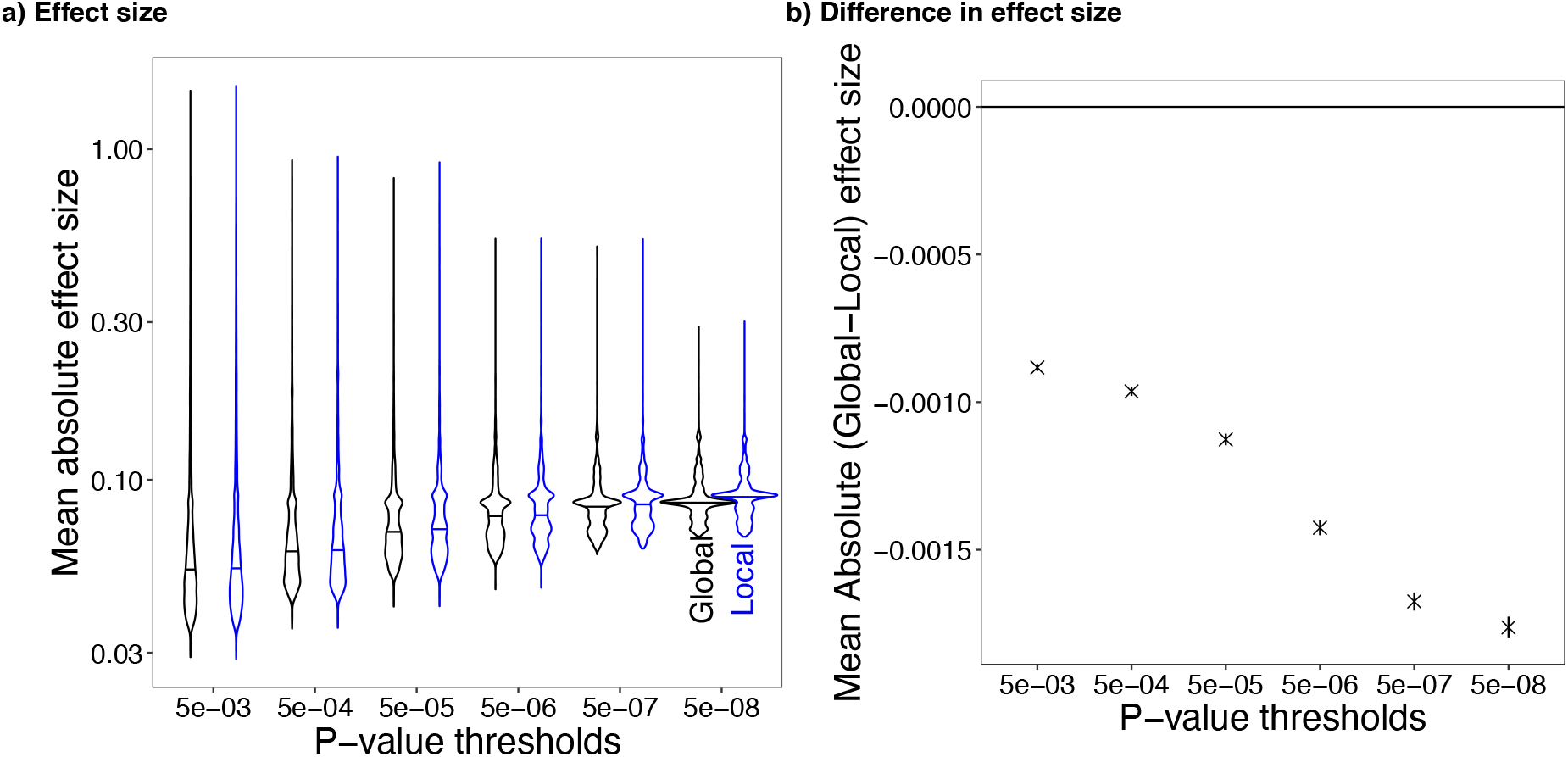
Stratification of SNP effect size bias in UK Biobank (education years) Stratification correction changes UK Biobank effect size estimates and the magnitude of the change varies by significance threshold. a) The mean absolute effect size for educational years and its median value as a function of p-value threshold, for Global or Local PC corrected meta-analysis. b) Mean absolute difference in effect size (Global – Local) effect size.

These results are consistent with the proportion of variance in different phenotypes (e.g. education attainment and BMI) being larger when corrected for global PCs than local PCs (Supplementary Figure S7). The magnitude of the difference varies between phenotypes, and depends on the causal model i.e. the relationships between phenotype, genotype, ancestry, and geography (Lawson et al. 2019).

### Reference PCs can be used to identify structure: a case study in ALSPAC

To test our hypothesis that uncorrected population structure may lead to misleading inference, we examined the ALSPAC cohort. Local variation is lost when effective sample size for a particular ancestry reduces beyond a threshold. We compare two studies in Bristol, the UK Biobank (N=27,503) and ALSPAC (N=7,927 mothers in our analysis). When constructing global ancestry using the entire UK Biobank variation, the two datasets have very similar genetic variation profiles across all PCs (Figure 4), including the main structures such as varying Scottish/English ancestry proportions.

**Figure 4.**
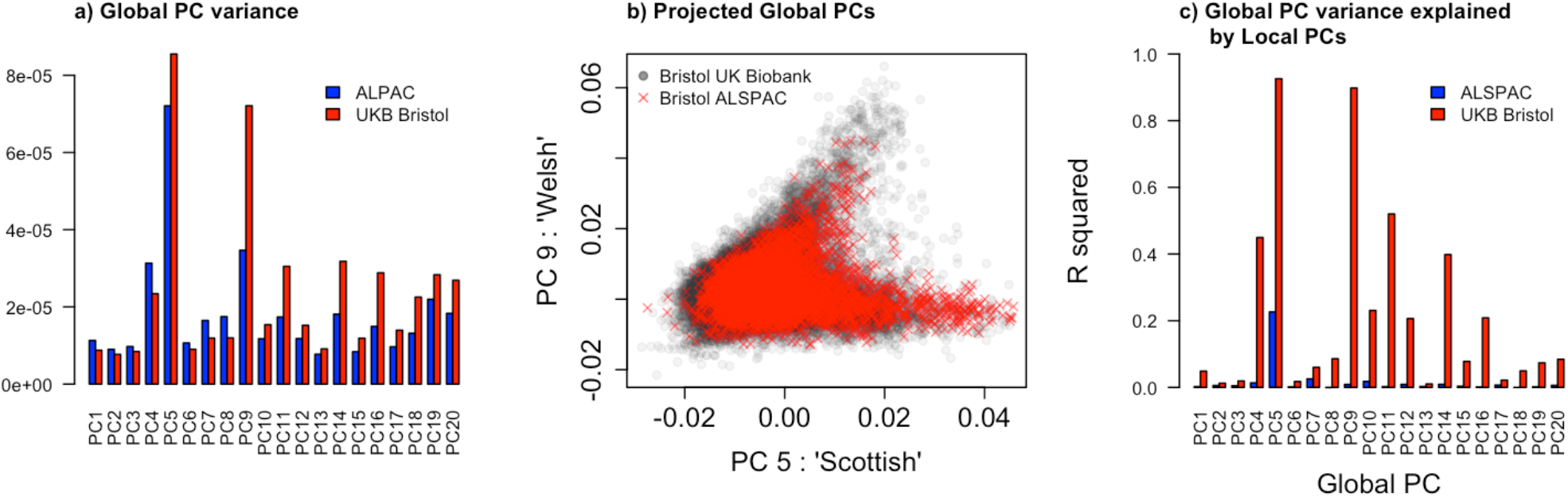
Population structure is lost in the ALSPAC cohort using local PCs. Local variation is lost when sample size reduces beyond a threshold as demonstrated by two studies in Bristol, the UK Biobank (N=27,503) and ALSPAC (N=7,927). a) Using global PCs constructed from UK Biobank variation, the two datasets have very similar genetic variation profiles across the first 20 global PCs. b) Comparing PC5 (high values associated with Scottish ancestry) and PC9 (high values associated with Welsh ancestry) the structure is similar. c) When projecting local PCs into global PCs, the proportion of variance explained is high for Bristol UK Biobank but very low within ALSPAC, due to sample size.

However, the datasets differ when projecting local ancestry PCs constructed from within each dataset into global ancestry (see Materials and Methods). Local PCs of the larger UK Biobank Bristol centre dataset partially recover most of the UK variation, whilst PCs of the smaller ALSPAC dataset recover very little. This would lead to systematic under-correction if replicated across a meta-analysis.

But does this matter for understanding phenotypes? To answer this question, we examined several phenotypes that have been studied with well-powered GWAS, including BMI, Educational attainment, IQ and C-reactive protein (CRP). We estimated the effect size in ALSPAC for both the study mothers and study children for SNPs identified by previous studies (see Materials and Methods) correcting either for local or global PCs.

Summarizing the total variance explained for phenotypes (Figure 5a) we find that the global PCs explain more variation in EA, IQ and BMI, but not CRP. This is most dramatic for mothers’ EA for which 7% vs 1% (global vs local) of variation is explained, matching previous estimates using haplotype information (Lawson et al. 2012) to quantify population structure in ALSPAC (Haworth et al. 2019; Lawson et al. 2020).

**Figure 5.**
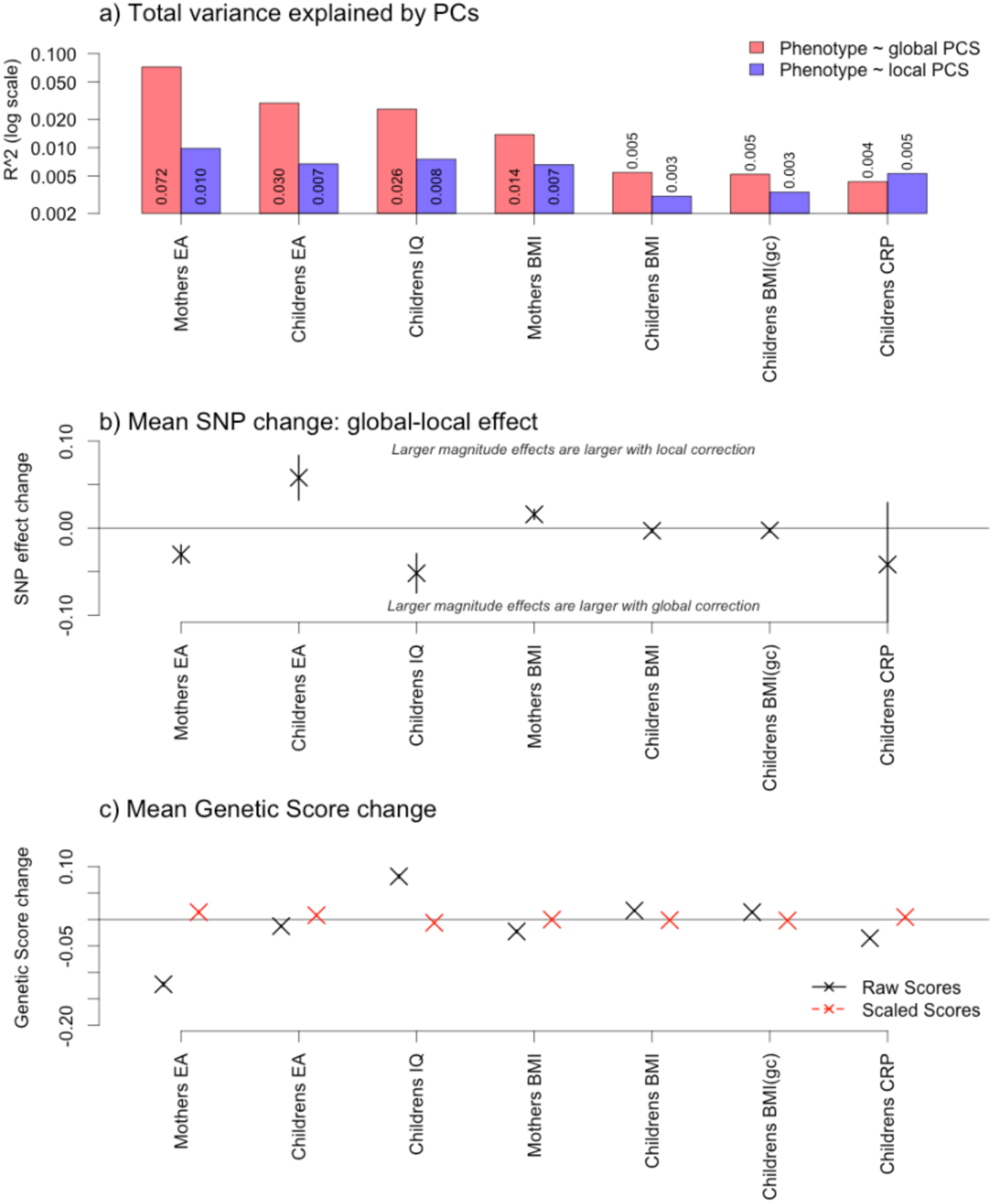
Stratification correction affects SNP inference in the ALSPAC cohort. Stratification correction choice makes a measurable impact on inferences from the ALSPAC cohort. a) Total variance in phenotype explained by global or local PCs (log scale). b-c) Weighted linear regression coefficients for measuring local PC bias. The regression coefficient 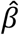 (and 95% confidence interval) from *δ*_*i*_ = *z*_*i,global*_ *− z*_*i,local*_ = *α* + *βz*_*i*_ + *ε*_*i*_, with *z*_*i*_ = (*z*_*i,global*_ + *z*_*i,local*_)/2 and in b) *z*_*i*_ is the SNP effect size for each GWAS. In c) *z*_*i*_ is the individual’s Genetic Score, either raw (summing the effect of each SNP present in the individual) or scaled to have mean 0 and s.d. 1 independently for both GWAS.

As ALSPAC is a relatively small cohort, the uncertainty involved in SNP effect estimation dominates the results. However, we found that the more robust estimates (higher Z-scores) changed systematically between correction models (Figure 5b, Supplementary Figure S8). Intriguingly, the direction is not the same for all phenotypes; local correction results in relatively larger estimates (i.e. under-correction) for EA, whilst it results in smaller estimates for BMI, which could imply subtle relatedness or improved power from correcting for ancestry.

Constructing a Genetic Score using this procedure leads to a similar picture, with systematic biases in prediction (Figure 5c, Supplementary Figure S9). Whilst there is statistical power to detect some differences in the scaled scores (e.g. in EA and CRP) these are unlikely to be practically significant changes. We therefore view the ordering of individuals to have been robust in this example. However, the raw scores are strongly skewed, again with biases in both directions, and further, the bias direction appears unrelated to whether SNPs were individually over or under predicted.

### Providing an appropriate set of ancestry covariates

The primary barrier to using the UK Biobank PCs is a lack of access to a) SNP loadings, and b) reference information to scale SNPs and perform QC carefully. We provide the key 18 ancestry PCs plus SNP information in an R package and script (see Code Availability) which allows trivial access to for all datasets in plink bim/bed/fam format of any size (e.g. runs on all 500k UK Biobank individuals in 6 hours). We further provide up to 200 UK Biobank PCs.

Users with access to UK Biobank data should consider the *bigsnpr* R package (Privé et al. 2020) which allows translation of any dataset into UK Biobank PCs with careful quality control assured due to comparison with the original raw data. Advanced users who do their own quality control and imputation may wish to directly apply the *flashpca* software (Abraham, Qiu, and Inouye 2017) to our provided reference data. Our package provides strand and build checks, automatically merges data coded with different minor alleles, and accounts for a moderate amount of non-overlapping SNPs.

Above, our UK Biobank results used BoltLMM (Loh et al. 2015). We confirm that these results are not meaningfully different to what we would have seen using linear regression correcting for PCs with PLINK (Supplementary Figure S10). The ALSPAC results also used PLINK. Therefore the effects describe are confirmed to apply to both linear regression and linear mixed models using the BoltLMM approximation.

## Discussion

Population stratification in association studies has received much attention. However, it has typically been considered as a problem of unintended correlations within the dataset, leading to correction in the form of a within-sample analysis (using PCA or other approaches). We provide evidence that this framing is insufficient. Whilst it is indeed unintended correlations that we wish to correct for, population structure is not always detectible from the dataset being studied. This hard-to-quantify population structure can be structurally related to phenotypes.

We demonstrated that within the UK Biobank’s individual study centres with samples of tens of thousands, as well as in the independent ALSPAC cohort, correcting for population stratification with a high-quality, external measure of population structure is necessary. Population structure exists at the within-city level and it is not correctly quantified within geographically clustered datasets. We found considerable residual correlation with phenotypes and identified that the SNP-level estimates were systematically biased. This resulted in appreciable error at the genome-wide level for the construction of Genetic Scores.

We identified that, were the UK Biobank to have been analysed as independent study centres subject to meta-analysis, then Educational Attainment, BMI, Psoriasis, Hyperthyroidism and Hypothyroidism would all have led to biased inference. This is likely to be the tip of the iceberg in meta-analyses, since the UK is a rather homogeneous population and the power in rare diseases is low.

Because global PCs are unarguably a better measure of population structure, it is tempting to interpret the effect size for the global PC correction as “more correct” than that for the local PC correction, and therefore the difference as a bias with the traditional approach. However, it is not that simple. We found little consistency in the direction of the bias; for example, EA for ALSPAC children appears to be “undercorrected” by local PCs, whereas the mothers EA appeared “overcorrected”. The reality is that confounding is caused by many sources, and shared ancestry is just one. Here we suspect that cryptic relatedness may exist, which is captured only by the local PCs.

The informed reader may find these results self-evident. However, the evidence that we provide of the importance and ease of improved stratification correction has clear implications. Future meta-analyses and association studies should adopt a new protocol for quantifying population stratification. Further, every analysis of small to medium sized cohorts whose association outputs remain of value should be re-considered. Large meta-analyses are particularly valuable and yet vulnerable to the biases identified here. Similarly, phenotypes with a non-trivial social or environmental component (Morris et al. 2020) are likely to be influenced by this or other hidden structural biases.

The new protocol should continue to adjust for relatedness within the cohort, but it must also add the confounding covariates of ancestry as quantified by a large and hence statistically powerful external resource. We provide such “genetic measures” for the UK Biobank reference in the form of PC loadings that can project any individual into this worldwide quantification of genetic variation.

Yet for non-UK individuals, even in the UK Biobank, this may be insufficient. There is no reason that institutions with access to large limited-access databases could not make and share independent PC loadings, for every region of the world, that smaller association studies with less power individually can apply. Although this is a partial solution because a nuanced quantification of ancestry is not linear, these sharable PCs will improve stratification correction with trivial cost, so the genetics community can and should implement this immediately.

## Data and Code availability

github.com/danjlawson/pcapred: R package for projecting into UK Biobank variation. github.com/danjlawson/pcapred-script: Script for non-R users to perform command line projection. github.com/danjlawson/pcapred-data: 200 ancestry PCs for UK Biobank.

ALSPAC (www.bristol.ac.uk/alspac/researchers/access/) and UK Biobank data (www.ukbiobank.ac.uk/principles-of-access/) are both accessible under their respective data use policies.

## Materials and Methods

### Cohorts

#### UK Biobank

The UK Biobank is a population-based health research resource consisting of approximately 500,000 people, aged between 38 years and 73 years, who were recruited between the years 2006 and 2010 from across the UK (Sudlow et al. 2015), particularly focused on identifying determinants of human diseases in middle-aged and older individuals, participants provided a range of information (such as demographics, health status, lifestyle measures, cognitive testing, personality self-report, and physical and mental health measures) via questionnaires and interviews; anthropometric measures, BP readings and samples of blood, urine and saliva were also taken (data available at www.ukbiobank.ac.uk). A full description of the study design, participants and quality control (QC) methods have been described in detail previously (Bycroft et al. 2018). UK Biobank received ethical approval from the Research Ethics Committee (REC reference for UK Biobank is 11/NW/0382). Access was under Application ID 21829.

#### ALSPAC

Pregnant women resident in Avon, UK with expected dates of delivery 1st April 1991 to 31st December 1992 were invited to take part in the study. The initial number of pregnancies enrolled is 14,541. Of these initial pregnancies, there was a total of 14,676 foetuses, resulting in 14,062 live births and 13,988 children who were alive at 1 year of age. When the oldest children were approximately 7 years of age, an attempt was made to bolster the initial sample with eligible cases who had failed to join the study originally. The total sample size for analyses using any data collected after the age of seven is therefore 15,454 pregnancies, resulting in 15,589 foetuses. Of these 14,901 were alive at 1 year of age. Ethical approval for the study was obtained from the ALSPAC Ethics and Law Committee and the Local Research Ethics Committees. Consent for biological samples has been collected in accordance with the Human Tissue Act (2004). Informed consent for the use of data collected via questionnaires and clinics was obtained from participants following the recommendations of the ALSPAC Ethics and Law Committee at the time. For further details of the cohort please see (Boyd et al. 2013; Fraser et al. 2013). Please note that the study website contains details of all the data that is available through a fullysearchabledatadictionaryandvariablesearchtool (http://www.bristol.ac.uk/alspac/researchers/our-data/).

### Genotyping, imputation and quality control

#### PCA Analysis

PCA analysis of the UK biobank was performed with *flashPCA* (Abraham, Qiu, and Inouye 2017) after following the procedure described in (Bycroft et al. 2018) to subset SNPs (147604 retained) and individuals (406758 retained). *FlashPCA* reports standardized Eigenvectors, unlike *FastPCA* (Galinsky et al. 2016) as used and reported by UK Biobank which scales Eigenvectors using the Eigenvalues. For stratification correction the distinction is not important, and our tool *pcapred* can translate between the two.

#### UK Biobank

The full data release contains the cohort of successfully genotyped samples (n=488,377). 49,979 individuals were genotyped using the UK BiLEVE array and 438,398 using the UK Biobank axiom array. Pre-imputation QC, phasing and imputation are described elsewhere (Bycroft et al. 2018). In brief, prior to phasing, multiallelic SNPs or those with MAF ≤1% were removed. Phasing of genotype data was performed using a modified version of the SHAPEIT2 algorithm (O’Connell et al. 2016). Genotype imputation to a reference set combining the UK10K haplotype and HRC reference panels 8was performed using IMPUTE2 algorithms (Howie, Marchini, and Stephens 2011). The analyses presented here were restricted to autosomal variants within the HRC site list using a graded filtering with varying imputation quality for different allele frequency ranges. Therefore, rarer genetic variants are required to have a higher imputation INFO score (Info>0.3 for MAF >3%; Info>0.6 for MAF 1-3%; Info>0.8 for MAF 0.5-1%; Info>0.9 for MAF 0.1-0.5%) with MAF and Info scores having been recalculated on an in-house derived ‘European’ subset (Mitchell et al. 2019).

Individuals with sex-mismatch (derived by comparing genetic sex and reported sex) or individuals with sex-chromosome aneuploidy were excluded from the analysis (n=814).

We restricted the sample to individuals of ‘european’ ancestry as defined by an in-house k-means cluster analysis performed using the first 4 principal components provided by UK Biobank in the statistical software environment R. The current analysis includes the largest cluster from this analysis (n=464,708) (Mitchell et al. 2019).

#### ALSPAC

DNA of the ALSPAC children was extracted from blood, cell line and mouthwash samples, then genotyped using references panels and subjected to standard quality control approaches. ALSPAC children were genotyped using the Illumina HumanHap550 quad chip genotyping platforms by 23andme subcontracting the Wellcome Trust Sanger Institute, Cambridge, UK and the Laboratory Corporation of America, Burlington, NC, US. The resulting raw genome-wide data were subjected to standard quality control methods. Individuals were excluded on the basis of gender mismatches; minimal or excessive heterozygosity; disproportionate levels of individual missingness (>3%) and insufficient sample replication (< 0.8). Population stratification was assessed by multidimensional scaling analysis and compared with Hapmap II (release 22) European descent (CEU), Han Chinese, Japanese and Yoruba reference populations; all individuals with non-European ancestry were removed. SNPs with a minor allele frequency of < 1%, a call rate of < 95% or evidence for violations of Hardy-Weinberg equilibrium (P < 5×10-7) were removed. Cryptic relatedness was measured as proportion of identity by descent (IBD) > 0.1. Related subjects that passed all other quality control thresholds were retained during subsequent phasing and imputation. 9,115 participants and 500,527 SNPs passed these quality control filters. ALSPAC mothers were genotyped using the Illumina human660W-quad array at Centre National de Génotypage (CNG) and genotypes were called with Illumina GenomeStudio. PLINK (v1.07) was used to carry out quality control measures on an initial set of 10,015 subjects and 557,124 directly genotyped SNPs. SNPs were removed if they displayed more than 5% missingness or a Hardy-Weinberg equilibrium P value of less than 1.0e-06. Additionally, SNPs with a minor allele frequency of less than 1% were removed. Samples were excluded if they displayed more than 5% missingness, had indeterminate X chromosome heterozygosity or extreme autosomal heterozygosity. Samples showing evidence of population stratification were identified by multidimensional scaling of genome-wide identity by state pairwise distances using the four HapMap populations as a reference, and then excluded. Cryptic relatedness was assessed using an IBD estimate of more than 0.125 which is expected to correspond to roughly 12.5% alleles shared IBD or a relatedness at the first cousin level. Related subjects that passed all other quality control thresholds were retained during subsequent phasing and imputation. 9,048 subjects and 526,688 SNPs passed these quality control filters.

We combined 477,482 SNP genotypes in common between the sample of mothers and sample of children. We removed SNPs with genotype missingness above 1% due to poor quality (11,396 SNPs removed) and removed a further 321 subjects due to potential ID mismatches. This resulted in a dataset of 17,842 subjects containing 6,305 duos and 465,740 SNPs (112 were removed during liftover and 234 were out of HWE after combination). We estimated haplotypes using ShapeIT (v2.r644) which utilises relatedness during phasing. The phased haplotypes were then imputed to the Haplotype Reference Consortium (HRCr1.1, 2016) panel of approximately 31,000 phased whole genomes. The HRC panel was phased using ShapeIt v2, and the imputation was performed using the Michigan imputation server. This gave 8,237 eligible children and 8,196 eligible mothers with available genotype data after exclusion of related subjects using cryptic relatedness measures described previously. Principal components were generated by extracting unrelated individuals (IBS < 0.05) and independent SNPs with long range LD regions removed, and then calculating using the ‘--pca’ command in plink1.90.

#### Association analysis: statistical methods

Genome-wide association analysis (GWAS) was conducted using linear mixed model (LMM) association method as implemented in BOLT-LMM (v2.3) (Loh et al. 2015). To model population structure in the sample we used 143,006 directly genotyped SNPs, obtained after filtering on MAF > 0.01; genotyping rate > 0.015; Hardy-Weinberg equilibrium p-value < 0.0001 and LD pruning to an r2 threshold of 0.1 using PLINKv2.00. Genotype array and sex were adjusted for in the model. BOLT-LMM association statistics are on the linear scale. As such, test statistics (betas and their corresponding standard errors) were transformed to log odds ratios and their corresponding 95% confidence intervals on the liability scale using a Taylor transformation expansion series (Loh et al. 2015).

#### Meta-analysis

Meta-analysis for variance explained was conducted using *rma* from the “metafor” package for R (Viechtbauer 2010) using the normal distribution approximation. P-values for the difference in *R*^*2*^ were calculated by computing the difference in the estimates, and the variance of the difference (the sum of the individual variances) and using the null that the *R*^*2*^*=0* again using *rma*. For binary traits, only study centres with at least 20 cases were considered. We also implemented a bootstrap procedure that did not make the normal distribution approximation, in which study centres were resampled 500 times with replacement. However, the results were not qualitatively different (not shown).

### Polygenic scoring

Genetic scores were created in the ALPAC cohort using PLINK (Purcell et al. 2007) based upon the list of SNPs identified to associate with educational attainment (Lee et al. 2018), BMI (Yengo et al. 2018), IQ (Lee et al. 2018) and CRP (Ligthart et al. 2018) in previous GWAS. All SNPs were weighted by their effect size in the replication cohort of the GWAS, and these sizes were summed using allelic scoring. The genetic scores were generated using GWAS results which had removed the ALSPAC cohort where included in the original GWAS, and therefore the scores are not perfectly comparable to those reported in the published meta-analysis. Where the lead SNPs from GWAS were not available in ALSPAC, we instead used the SNPs in highest linkage disequilibrium. Genetic score analysis in ALSPAC was run on age and sex standardised phenotypes controlling for either local PCs (the first 20 principal components of ancestry as identified within the ALSPAC cohort) or global PCs (the first 20 principal components of ancestry constructed from UK Biobank loadings).

#### ALSPAC phenotypes

For ALSPAC mothers, years of education was determined by recoding highest level of education reported during pregnancy. Response were coded as basic formal education (7 years), certificate of secondary education (10 years), O-levels and vocational qualifications (11 years), A-level (13 years), and degree (16 years). Mother’s BMI was measured during the ‘Focus on Mothers 1’ direct assessment when the study offspring were aged 17 (mother ages 34-63).

For ALSPAC children, education was measured using average fine graded point scores in age 16 educational examinations, which represents final compulsory schooling examinations. Scores were obtained through data linkage to the UK National Pupil Database (NPD), which represents the most accurate record of individual educational achievement available in the UK. Intelligence was measured during the direct assessment at age eight using the short form Wechsler Intelligence Scale for Children (WISC) (Wechsler 1992) from verbal, performance, and digit span tests and administered by members of the ALSPAC psychology team overseen by an expert in psychometric testing. Raw scores were recalculated to be comparable to those that would have been obtained had the full test been administered and then age-scaled to give a total overall score combined from the performance and verbal subscales. BMI was measured during the direct assessments at ages 7, 8, 9, 10 and 11. In order to increase sample size, where BMI data were not available at age 7 we used BMI measured at the earliest available subsequent measurement. C-reactive protein (CRP) was measured from non-fasting blood assays taken during direct assessment when the offspring were aged 9.

### Detecting bias in scores and SNP effects

To assess statistical power, we work with z-scores, i.e. 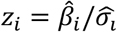, where 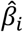 is the estimate of the effect of SNP *I* and 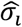 is the estimate of the standard deviation of this estimate. To compare the global (g) and local (l) effects we consider the mean estimate 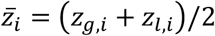 and difference *∂*_*i*_ = (*z*_*g,i*_ *− z*_*l,i*_) for each SNP *i*. To prevent the large number of barely-significant estimates from dominating the signal, we assign a weight to each SNP *w*_*i*_ = 1/*ρ*_*i*_ where *ρ*_*i*_ is the density estimate taken from a 5 nearest-neighbour estimate using “knnDE” from the R Package “TDA”. We then perform robust regression for *∂*∼*z* and report the regression estimate and confidence interval. We further checked that our conclusions are not impacted by these choices by performing regular unweighted regression for *∂*∼*z*.

### UK Biobank trait definition

Years of education was determined by recoding highest level of education reported in a questionnaire. Response were coded as basic formal education (7 years), O-levels/GCSEs/CSEs or equivalent (10 years), A-level/AS levels or equivalent (13 years), NVQ or HND or HNC or equivalent (19 years) and College/University degree (20 years). We also binary studied educational attainment (EA), which is measured as 1 for people who have obtained a College or University degree.

Height and weight were measured during the participants’ baseline visit to a UK Biobank assessment center.

Heel bone mineral density (eBMD) was estimated based on an ultrasound measurement of the calcaneus by UK Biobank. The T-score is the number of standard deviations for bone mineral density relative to the mean. Consistent with the criteria established by Kemp et al., individuals were excluded that exceeded the following thresholds for eBMD: males, ≤0.18 or ≥1.06 g/cm2; females ≤0.12 or ≥1.025 g/cm2.

Other traits were self-reported at the verbal interview and coded as yes/no. If the participant was uncertain of the type of illness they had had, then they described it to the interviewer (a trained nurse) who attempted to place it within the coding tree. If the illness could not be located in the coding tree then the interviewer entered a free-text description of it. These free-text descriptions were subsequently examined by a doctor and, where possible, matched to entries in the coding tree.

## Acknowledgements

We are extremely grateful to all the families who took part in this study, the midwives for their help in recruiting them, and the whole ALSPAC team, which includes interviewers, computer and laboratory technicians, clerical workers, research scientists, volunteers, managers, receptionists and nurses.

## Bibliography

Abdellaoui, Abdel, David Hugh-Jones, Loic Yengo, Kathryn E. Kemper, Michel G. Nivard, Laura Veul, Yan Holtz, et al. 2019. ‘Genetic Correlates of Social Stratification in Great Britain’. Nature Human Behaviour 3 (12): 1332–42. https://doi.org/10.1038/s41562-019-0757-5.

Abraham, Gad, Yixuan Qiu, and Michael Inouye. 2017. ‘FlashPCA2: Principal Component Analysis of Biobank-Scale Genotype Datasets’. Bioinformatics 33 (17): 2776–78. https://doi.org/10.1093/bioinformatics/btx299.

Berg, Jeremy J, Arbel Harpak, Nasa Sinnott-Armstrong, Anja Moltke Joergensen, Hakhamanesh Mostafavi, Yair Field, Evan August Boyle, et al. 2019. ‘Reduced Signal for Polygenic Adaptation of Height in UK Biobank’. Edited by Magnus Nordborg, Mark I McCarthy, Magnus Nordborg, Nicholas H Barton, and Joachim Hermisson. ELife 8 (March): e39725. https://doi.org/10.7554/eLife.39725.

Boyd, Andy, Jean Golding, John Macleod, Debbie A Lawlor, Abigail Fraser, John Henderson, Lynn Molloy, Andy Ness, Susan Ring, and George Davey Smith. 2013. ‘Cohort Profile: The “Children of the 90s”—the Index Offspring of the Avon Longitudinal Study of Parents and Children’. International Journal of Epidemiology 42 (1): 111–27. https://doi.org/10.1093/ije/dys064.

Bush, William S., and Jason H. Moore. 2012. ‘Chapter 11: Genome-Wide Association Studies’. PLoS Computational Biology 8 (12): e1002822. https://doi.org/10.1371/journal.pcbi.1002822.

Bycroft, Clare, Colin Freeman, Desislava Petkova, Gavin Band, Lloyd T. Elliott, Kevin Sharp, Allan Motyer, et al. 2018. ‘The UK Biobank Resource with Deep Phenotyping and Genomic Data’. Nature 562 (7726): 203–9. https://doi.org/10.1038/s41586-018-0579-z.

Evangelou, Evangelos, and John P. A. Ioannidis. 2013. ‘Meta-Analysis Methods for Genome-Wide Association Studies and Beyond’. Nature Reviews. Genetics 14 (6): 379–89. https://doi.org/10.1038/nrg3472.

Fraser, Abigail, Corrie Macdonald-Wallis, Kate Tilling, Andy Boyd, Jean Golding, George Davey Smith, John Henderson, et al. 2013. ‘Cohort Profile: The Avon Longitudinal Study of Parents and Children: ALSPAC Mothers Cohort’. International Journal of Epidemiology 42 (1): 97–110. https://doi.org/10.1093/ije/dys066.

Galinsky, Kevin J., Gaurav Bhatia, Po-Ru Loh, Stoyan Georgiev, Sayan Mukherjee, Nick J. Patterson, and Alkes L. Price. 2016. ‘Fast Principal-Component Analysis Reveals Convergent Evolution of ADH1B in Europe and East Asia’. The American Journal of Human Genetics 98 (3): 456–72. https://doi.org/10.1016/j.ajhg.2015.12.022.

GIANT Consortium. 2018. 2018. portals.broadinstitute.org/collaboration/giant/index.php/GIANT_consortium.

Haworth, Simon, Ruth Mitchell, Laura Corbin, Kaitlin H. Wade, Tom Dudding, Ashley Budu-Aggrey, David Carslake, et al. 2019. ‘Apparent Latent Structure within the UK Biobank Sample Has Implications for Epidemiological Analysis’. Nature Communications 10 (1): 333. https://doi.org/10.1038/s41467-018-08219-1.

Howie, Bryan, Jonathan Marchini, and Matthew Stephens. 2011. ‘Genotype Imputation with Thousands of Genomes’. G3: Genes|Genomes|Genetics 1 (6): 457–70. https://doi.org/10.1534/g3.111.001198.

Lawson, Daniel John, Neil Martin Davies, Simon Haworth, Bilal Ashraf, Laurence Howe, Andrew Crawford, Gibran Hemani, George Davey Smith, and Nicholas John Timpson. 2020. ‘Is Population Structure in the Genetic Biobank Era Irrelevant, a Challenge, or an Opportunity?’ Human Genetics 139 (1): 23–41. https://doi.org/10.1007/s00439-019-02014-8.

Lawson, Daniel John, and Daniel Falush. 2012. ‘Population Identification Using Genetic Data’. Annual Review of Genomics and Human Genetics 13 (1): 337–61. https://doi.org/10.1146/annurev-genom-082410-101510.

Lawson, Daniel John, Garrett Hellenthal, Simon Myers, and Daniel Falush. 2012. ‘Inference of Population Structure Using Dense Haplotype Data’. PLOS Genet 8 (1): e1002453. https://doi.org/10.1371/journal.pgen.1002453.

Lee, James J., Robbee Wedow, Aysu Okbay, Edward Kong, Omeed Maghzian, Meghan Zacher, Tuan Anh Nguyen-Viet, et al. 2018. ‘Gene Discovery and Polygenic Prediction from a Genome-Wide Association Study of Educational Attainment in 1.1 Million Individuals’. Nature Genetics 50 (8): 1112–21. https://doi.org/10.1038/s41588-018-0147-3.

Leslie, Stephen, Bruce Winney, Garrett Hellenthal, Dan Davison, Abdelhamid Boumertit, Tammy Day, Katarzyna Hutnik, et al. 2015. ‘The Fine-Scale Genetic Structure of the British Population’. Nature 519 (7543): 309–14. https://doi.org/10.1038/nature14230.

Li, Yun, Cristen Willer, Serena Sanna, and Gonçalo Abecasis. 2009. ‘Genotype Imputation’. Annual Review of Genomics and Human Genetics 10: 387–406. https://doi.org/10.1146/annurev.genom.9.081307.164242.

Ligthart, Symen, Ahmad Vaez, Urmo Võsa, Maria G. Stathopoulou, Paul S. de Vries, Bram P. Prins, Peter J. Van der Most, et al. 2018. ‘Genome Analyses of >200,000 Individuals Identify 58 Loci for Chronic Inflammation and Highlight Pathways That Link Inflammation and Complex Disorders’. The American Journal of Human Genetics 103 (5): 691–706. https://doi.org/10.1016/j.ajhg.2018.09.009.

Linnér, Richard K., Pietro Biroli, Edward Kong, S. Fleur W. Meddens, Robbee Wedow, Mark Alan Fontana, Maël Lebreton, et al. 2019. ‘Genome-Wide Association Analyses of Risk Tolerance and Risky Behaviors in over 1 Million Individuals Identify Hundreds of Loci and Shared Genetic Influences’. Nature Genetics 51 (2): 245–57. https://doi.org/10.1038/s41588-018-0309-3.

Locke, Adam E., Bratati Kahali, Sonja I. Berndt, Anne E. Justice, Tune H. Pers, Felix R. Day, Corey Powell, et al. 2015. ‘Genetic Studies of Body Mass Index Yield New Insights for Obesity Biology’. Nature 518 (7538): 197–206. https://doi.org/10.1038/nature14177.

Loh, Po-Ru, George Tucker, Brendan K. Bulik-Sullivan, Bjarni J. Vilhjálmsson, Hilary K. Finucane, Rany M. Salem, Daniel I. Chasman, et al. 2015. ‘Efficient Bayesian Mixed-Model Analysis Increases Association Power in Large Cohorts’. Nature Genetics 47 (3): 284–90. https://doi.org/10.1038/ng.3190.

Mitchell, Ruth, Gibran Hemani, Tom Dudding, Laura Corbin, Sean Harrison, and Lavinia Paternoster. 2019. ‘UK Biobank Genetic Data: MRC-IEU Quality Control, Version 2.’ https://doi.org/10.5523/bris.1ovaau5sxunp2cv8rcy88688v.

Morris, Tim T., Neil M. Davies, Gibran Hemani, and George Davey Smith. 2020. ‘Population Phenomena Inflate Genetic Associations of Complex Social Traits’. Science Advances 6 (16): eaay0328. https://doi.org/10.1126/sciadv.aay0328.

O’Connell, Jared, Kevin Sharp, Nick Shrine, Louise Wain, Ian Hall, Martin Tobin, Jean-Francois Zagury, Olivier Delaneau, and Jonathan Marchini. 2016. ‘Haplotype Estimation for Biobank Scale Datasets’. Nature Genetics 48 (7): 817–20. https://doi.org/10.1038/ng.3583.

Okbay, Aysu, Jonathan P. Beauchamp, Mark Alan Fontana, James J. Lee, Tune H. Pers, Cornelius A. Rietveld, Patrick Turley, et al. 2016. ‘Genome-Wide Association Study Identifies 74 Loci Associated with Educational Attainment’. Nature 533 (7604): 539–42. https://doi.org/10.1038/nature17671.

Paternoster, Lavinia, Kate Tilling, and George Davey Smith. 2017. ‘Genetic Epidemiology and Mendelian Randomization for Informing Disease Therapeutics: Conceptual and Methodological Challenges’. PLOS Genetics 13 (10): e1006944. https://doi.org/10.1371/journal.pgen.1006944.

Peterson, Roseann E., Alexis C. Edwards, Silviu-Alin Bacanu, Danielle M. Dick, Kenneth S. Kendler, and Bradley T. Webb. 2017. ‘The Utility of Empirically Assigning Ancestry Groups in Cross-Population Genetic Studies of Addiction’. The American Journal on Addictions 26 (5): 494–501. https://doi.org/10.1111/ajad.12586.

Privé, Florian, Keurcien Luu, Michael G. B. Blum, John J. McGrath, and Bjarni J. Vilhjálmsson. 2020. ‘Efficient Toolkit Implementing Best Practices for Principal Component Analysis of Population Genetic Data’. BioRxiv, January, 841452. https://doi.org/10.1101/841452.

Purcell, Shaun, Benjamin Neale, Kathe Todd-Brown, Lori Thomas, Manuel A. R. Ferreira, David Bender, Julian Maller, et al. 2007. ‘PLINK: A Tool Set for Whole-Genome Association and Population-Based Linkage Analyses’. American Journal of Human Genetics 81 (3): 559–75. https://doi.org/10.1086/519795.

Reisberg, Sulev, Tatjana Iljasenko, Kristi Läll, Krista Fischer, and Jaak Vilo. 2017. ‘Comparing Distributions of Polygenic Risk Scores of Type 2 Diabetes and Coronary Heart Disease within Different Populations’. PloS One 12 (7): e0179238. https://doi.org/10.1371/journal.pone.0179238.

Sohail, Mashaal, Robert M Maier, Andrea Ganna, Alex Bloemendal, Alicia R Martin, Michael C Turchin, Charleston WK Chiang, et al. 2019. ‘Polygenic Adaptation on Height Is Overestimated Due to Uncorrected Stratification in Genome-Wide Association Studies’. Edited by Magnus Nordborg, Mark I McCarthy, Magnus Nordborg, Nicholas H Barton, and Joachim Hermisson. ELife 8 (March): e39702. https://doi.org/10.7554/eLife.39702.

Sudlow, Cathie, John Gallacher, Naomi Allen, Valerie Beral, Paul Burton, John Danesh, Paul Downey, et al. 2015. ‘UK Biobank: An Open Access Resource for Identifying the Causes of a Wide Range of Complex Diseases of Middle and Old Age’. PLOS Medicine 12 (3): e1001779. https://doi.org/10.1371/journal.pmed.1001779.

Viechtbauer, Wolfgang. 2010. ‘Conducting Meta-Analyses in R with the Metafor Package’. Journal of Statistical Software 36 (3): 1–48. https://doi.org/10.18637/jss.v036.i03.

Visscher, Peter M., Naomi R. Wray, Qian Zhang, Pamela Sklar, Mark I. McCarthy, Matthew A. Brown, and Jian Yang. 2017. ‘10 Years of GWAS Discovery: Biology, Function, and Translation’. The American Journal of Human Genetics 101 (1): 5–22. https://doi.org/10.1016/j.ajhg.2017.06.005.

Warrington, Nicole M., Laura D. Howe, Lavinia Paternoster, Marika Kaakinen, Sauli Herrala, Ville Huikari, Yan Yan Wu, et al. 2015. ‘A Genome-Wide Association Study of Body Mass Index across Early Life and Childhood’. International Journal of Epidemiology 44 (2): 700–712. https://doi.org/10.1093/ije/dyv077.

Wechsler, D. 1992. Wechsler Intelligence Scle for Children. Third Edition. The Psychological Corporation.

Wellcome Trust Case Control Consortium. 2007. ‘Genome-Wide Association Study of 14,000 Cases of Seven Common Diseases and 3,000 Shared Controls’. Nature 447 (7145): 661–78. https://doi.org/10.1038/nature05911.

Wood, Andrew R., Tonu Esko, Jian Yang, Sailaja Vedantam, Tune H. Pers, Stefan Gustafsson, Audrey Y. Chu, et al. 2014. ‘Defining the Role of Common Variation in the Genomic and Biological Architecture of Adult Human Height’. Nature Genetics 46 (11): 1173–86. https://doi.org/10.1038/ng.3097.

Yang, Jian, S. Hong Lee, Michael E. Goddard, and Peter M. Visscher. 2011. ‘GCTA: A Tool for Genome-Wide Complex Trait Analysis’. The American Journal of Human Genetics 88 (1): 76–82. https://doi.org/10.1016/j.ajhg.2010.11.011.

Yengo, Loic, Julia Sidorenko, Kathryn E. Kemper, Zhili Zheng, Andrew R. Wood, Michael N. Weedon, Timothy M. Frayling, et al. 2018. ‘Meta-Analysis of Genome-Wide Association Studies for Height and Body Mass Index in ∼700,000 Individuals of European Ancestry’. BioRxiv, March, 274654. https://doi.org/10.1101/274654.

Yu, Jianming, Gael Pressoir, William H. Briggs, Irie Vroh Bi, Masanori Yamasaki, John F. Doebley, Michael D. McMullen, et al. 2006. ‘A Unified Mixed-Model Method for Association Mapping That Accounts for Multiple Levels of Relatedness’. Nature Genetics 38 (2): 203–8. https://doi.org/10.1038/ng1702.

Zeggini, Eleftheria, and John P. A. Ioannidis. 2009. ‘Meta-Analysis in Genome-Wide Association Studies’. Pharmacogenomics 10 (2): 191–201. https://doi.org/10.2217/14622416.10.2.191.

Zhu, Zhihong, Zhili Zheng, Futao Zhang, Yang Wu, Maciej Trzaskowski, Robert Maier, Matthew R. Robinson, et al. 2018. ‘Causal Associations between Risk Factors and Common Diseases Inferred from GWAS Summary Data’. Nature Communications 9 (1): 224. https://doi.org/10.1038/s41467-017-02317-2.

