## Supplementary Figures S1-10 for "Population stratification in GWAS meta-analysis should be standardized to the best available reference datasets"

#### Supplementary Figure S1: PC interpretation

LD Structure as a function of PC in UK Biobank. a) Average auto-correlation function (ACF) at lag 1 of absolute value of loadings, for all PCs. b) aggregated loadings from highlighted PCs (red in a) when SNPs are aggregated into bins of 100.

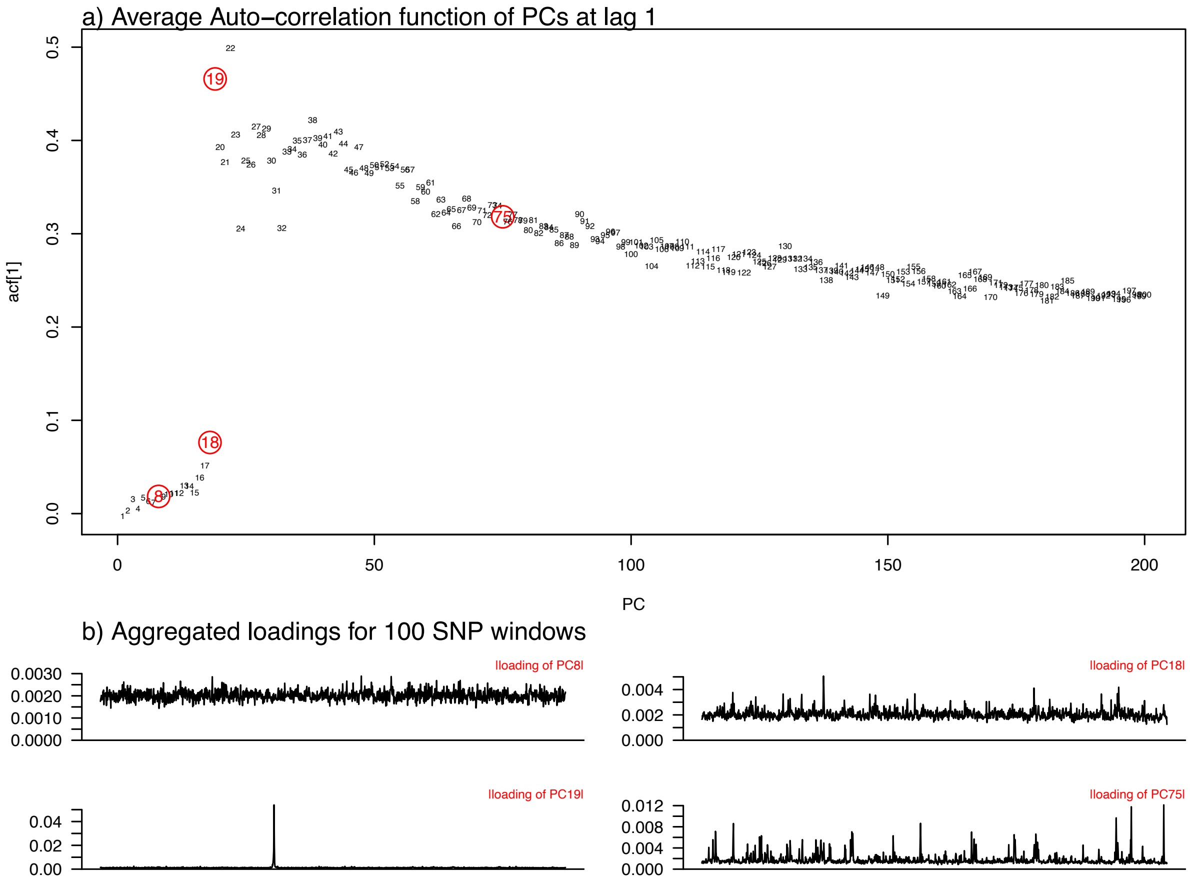

#### Supplementary Figure S2: Mean PC distribution by study centre

Coloured from minimum value (dark blue) to maximum value (light blue).

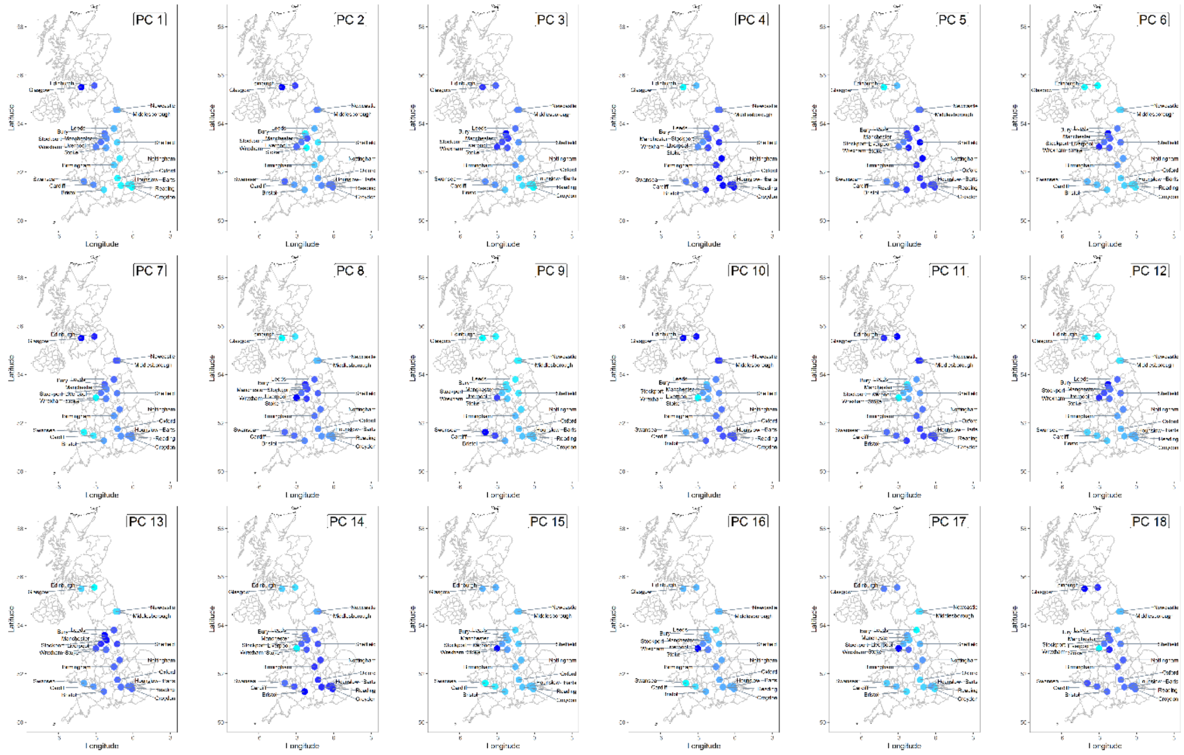


#### Supplementary Figure S3: Mean PCs by study centre

a) PC5 (x-axis) vs PC9 (y-axis). b) PC11 (x-axis) vs PC 14 (y-axis).

Colour gradient: from North-East (blue) to South-West (yellow) with red in the middle.

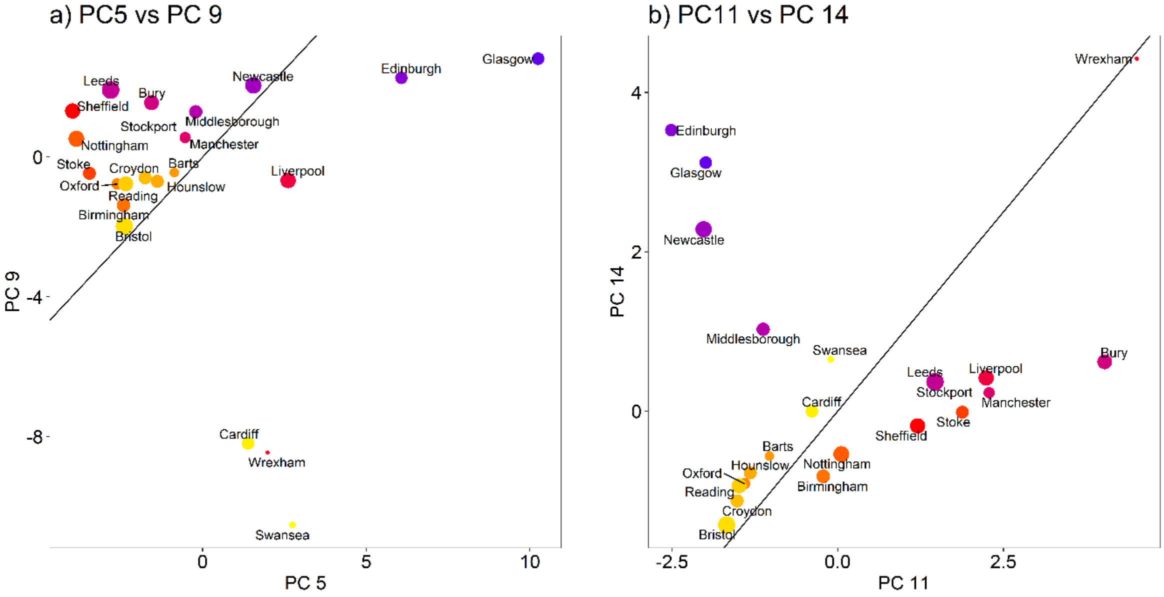

### Supplementary Figure S4. UK Biobank traits distribution by study centre

Summaries of phenotype variation by study centre for BMI, years of education, educational attainment. All traits vary systematically by geography.

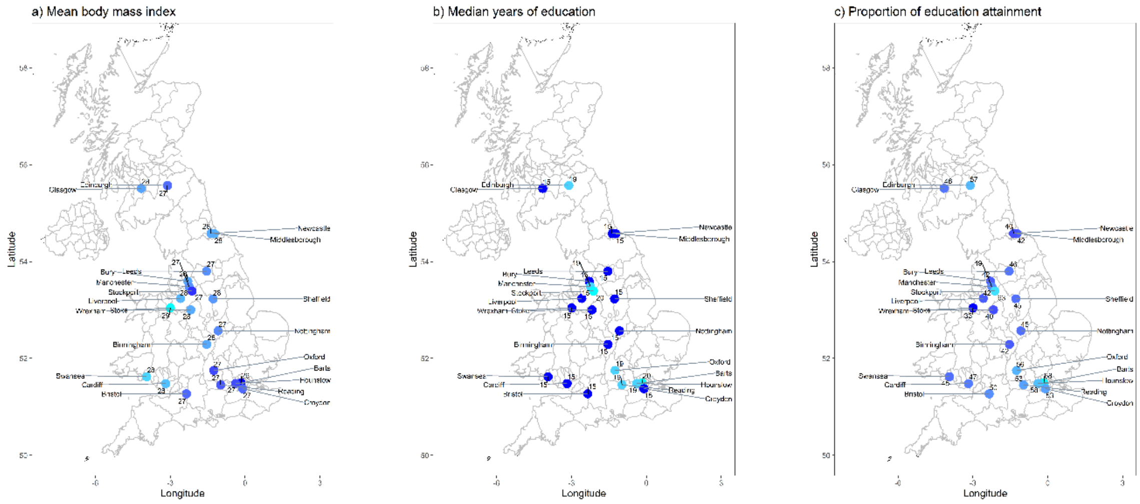

#### Supplementary Figure S5: UK Biobank disease traits

Proportion of variance explained in a meta-analysis for 24 binary disease outcomes, plus Bone mineral density and Height. Shown is the combined estimate from a meta analysis where each study centre is corrected either for Global (i.e. UK Biobank whole) or Local (i.e. study centre specific) Principal Components, as well as the p-value for the difference (see Methods), and the number of cases of the trait (which equals the sample size for continuous traits).

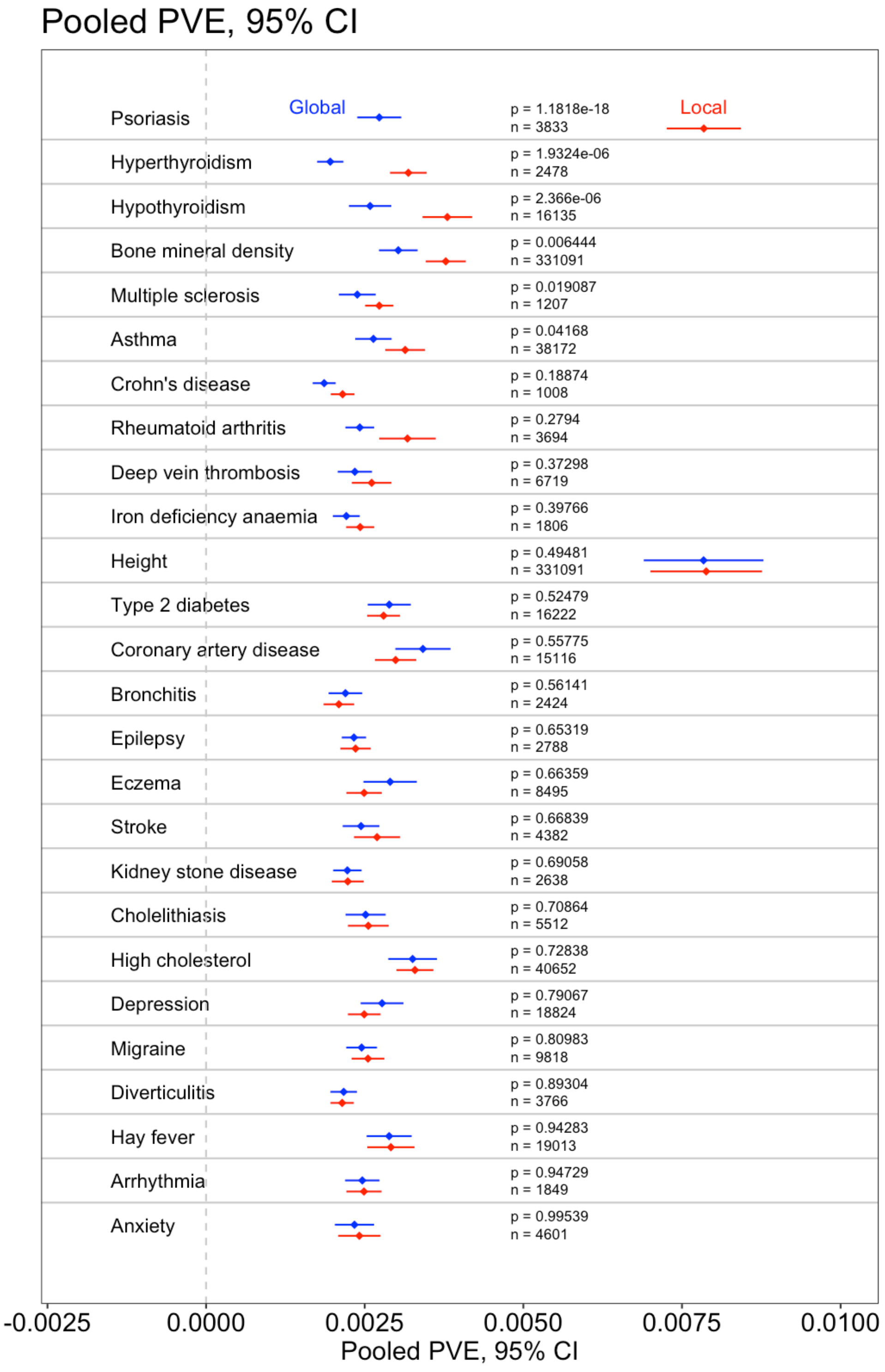

#### Supplementary Figure S6: UK Biobank SNP effect size by PC correction

Scatterplots of effect sizes in UK Biobank for Local vs Global PC correction, for a) EA, b) BMI, showing only SNPs significant in at least one correction. The slope of the unconstrained regression line shows is given in the title.

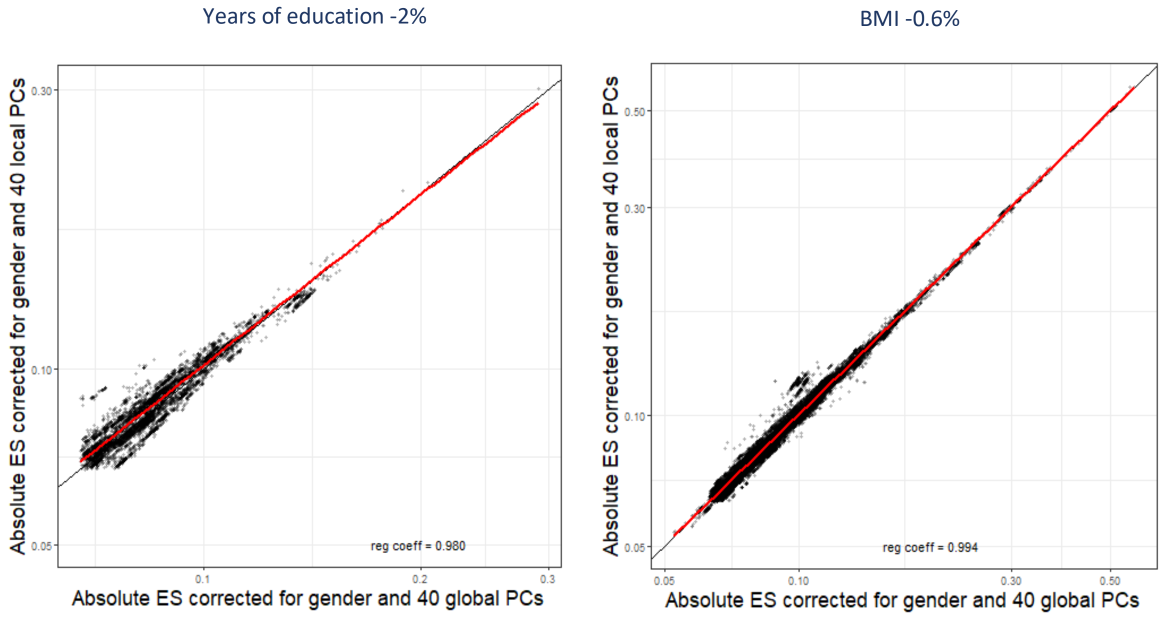

#### Supplementary Figure S7. Proportion of variance explained by local and global PCs

Proportion of Variance explained as a function of study centre size for a) in Educational attainment and b) BMI.

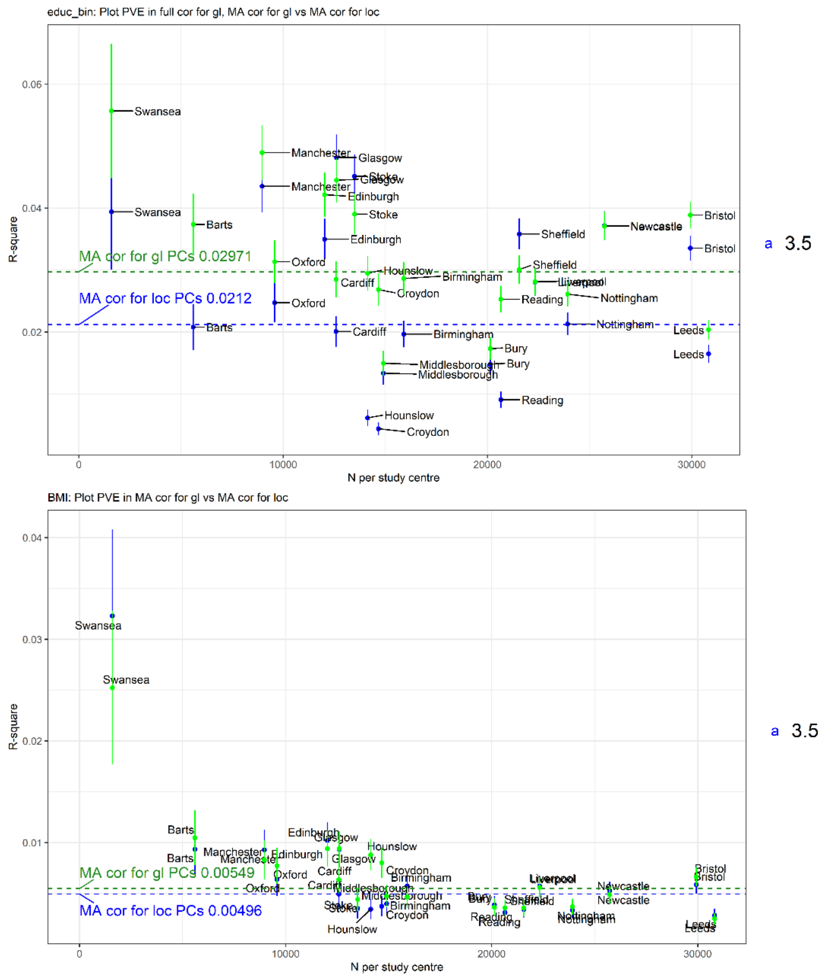


#### Supplementary Figure S8. Correction for individual SNP effects in ALSPAC

Effect size change as a function of effect magnitude. On the x-axis we show the average effect across global and local PC correction. On the y-axis we show the UKB effect size – ALSPAC effect size. Shown are two regression lines, standard and weighted inversely with sample size. Each subfigure a-g) shows a different phenotype/cohort pair.

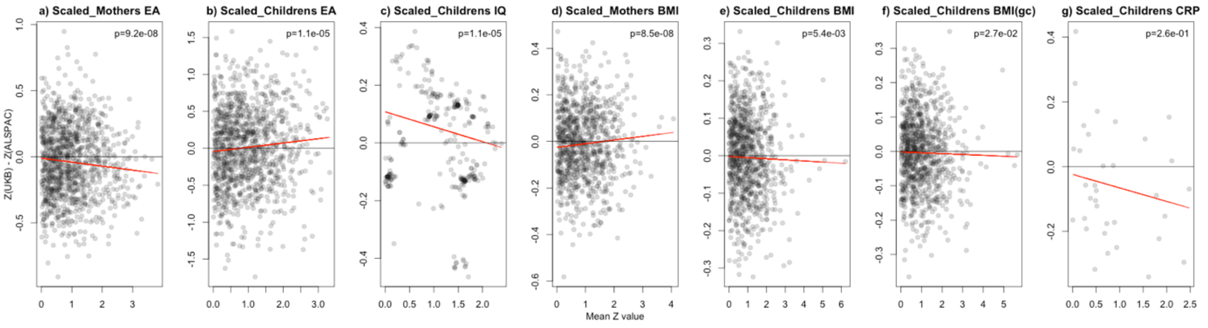

#### Supplementary Figure S9. Correction for Genetic Scores in ALSPAC

Effect size change as a function of effect magnitude. On the x-axis we show the average effect across global and local PC correction. On the y-axis we show the UKB effect size – ALSPAC effect size. Shown are two regression lines, standard and weighted inversely with sample size. a-g are for the raw genetic scores, h-n are for the standardized genetic scores.

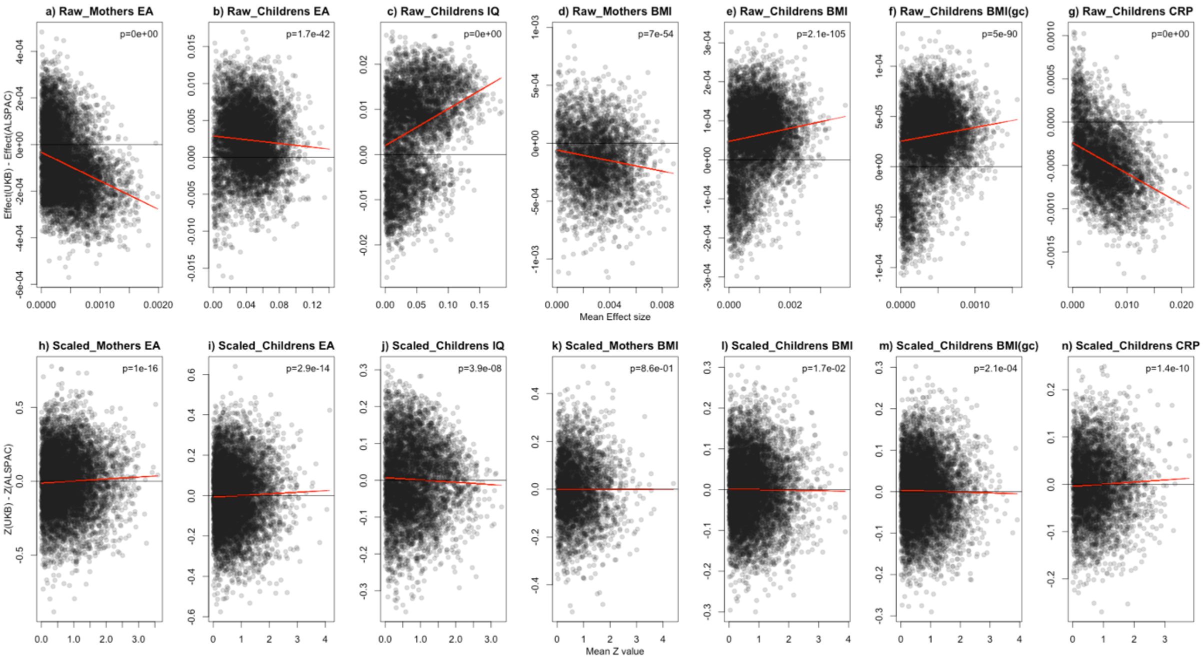


#### Supplementary Figure S10. Comparison of Bolt-LMM and PLINK Linear Regression for UK Biobank Educational Attainment

Effect size estimates are consistent between the Linear Mixed Model of BOLT-LMM vs the standard regression implemented in PLINK, performing meta-analysis using global PCs for education attainment binary outcome.

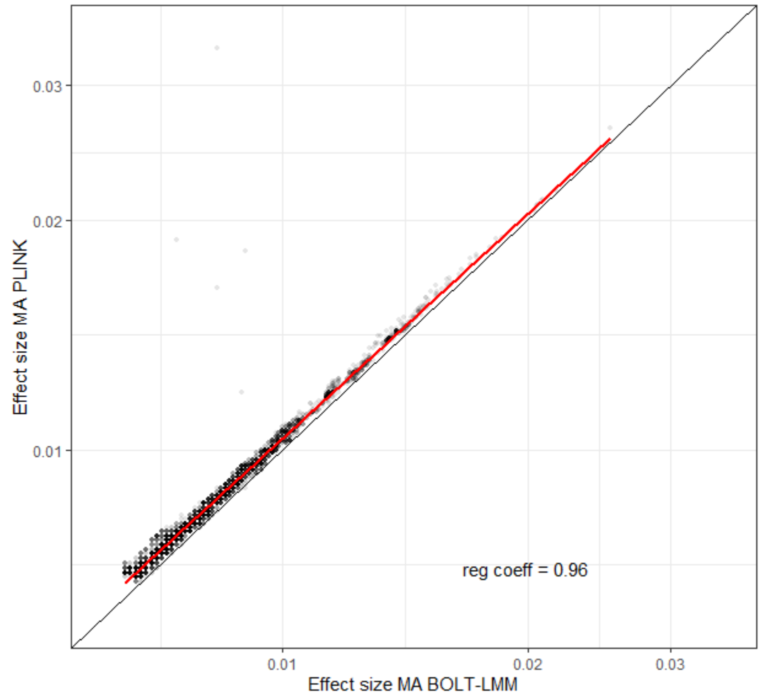
